# Self-other representation in the social brain reflects social connection

**DOI:** 10.1101/856856

**Authors:** Andrea L. Courtney, Meghan L. Meyer

## Abstract

Social connection is critical to well-being, yet how the brain reflects our attachment to other people remains largely unknown. We combined univariate and multivariate brain imaging analyses to assess whether and how the brain organizes representations of others based on how connected they are to our own identity. During an fMRI scan, participants (N=43) completed a self- and other-reflection task for 16 targets: the self, five close others, five acquaintances, and five celebrities. In addition, they reported their subjective closeness to each target and their own trait loneliness. We examined neural responses to the self and others in a brain region that has been associated with self-representation (medial prefrontal cortex; MPFC) and across the whole brain. The structure of self-other representation in the MPFC and across the social brain appeared to cluster targets into three social categories: the self, social network members (including close others and acquaintances), and celebrities. Moreover, both univariate activation in MPFC and multivariate self-other similarity in MPFC and across the social brain increased with subjective self-other closeness ratings. Critically, participants who were less socially connected (i.e. lonelier) showed altered self-other mapping in social brain regions. Most notably, in MPFC, loneliness was associated with reduced representational similarity between the self and others. The social brain apparently maintains information about broad social categories as well as closeness to the self. Moreover, these results point to the possibility that feelings of chronic social disconnection may be mirrored by a ‘lonelier’ neural self-representation.

**Significance Statement:** Social connection is critical to well-being, yet how the brain reflects our attachment to people remains unclear. We found that the social brain stratifies neural representations of people based on our subjective connection to them, separately clustering people who are and are not in our social network. Moreover, the people we feel closest to are represented most closely to ourselves. Finally, lonelier individuals also appeared to have a ‘lonelier’ neural self-representation in the MPFC, as loneliness attenuated the closeness between self and other neural representations in this region. The social brain appears to map our interpersonal ties, and alterations in this map may help explain why lonely individuals endorse statements such as ‘people are around me but not with me’.

## Introduction

Feeling close to other people promotes well-being (Barnett and Gotlib, 1988; Diener and Seligman, 2002; Holt-Lunstad et al., 2010) whereas feeling disconnected from them can compromise mental and physical health (Baumeister and Leary, 1995; Baumeister and Tice, 1990; Cacioppo et al., 2006). Yet, how the brain represents interpersonal closeness remains unclear. Filling this gap is critical as it may reveal basic mechanisms to intervene on to increase subjective connection and in turn promote well-being.

Insight into how the brain represents subjective social connection may come from a close examination of the medial prefrontal cortex (MPFC). While the MPFC is known to preferentially activate in response to thinking about the self, it exhibits similar activation when thinking about close others (Seger et al., 2004; Heatherton et al., 2006; Krienen et al., 2010; Moran et al., 2011; Chen et al., 2013). Moreover, these activation levels hold after controlling for one’s similarity to the close other considered (Krienen et al., 2010), and are elicited more strongly by deeper characteristics of the person (e.g., their personality) than by superficial characteristics (e.g., their appearance; Moran et al., 2011). Collectively, these results suggest that the MPFC may play a key role in representing our personal connection to others.

If the MPFC represents our social connection to others, there are at least two ways in which it may pull this off. One possibility is that the MPFC keeps a structured map of our social circles, with people organized by how close they are to us (Dunbar, 2018). Consistent with this possibility, MPFC responses reflect others’ *objective* social network positions (Parkinson et al., 2017). For example, multivariate MPFC responses to viewing people from one’s own social network mirror their eigenvector centrality (i.e., the extent to which they are connected to well-connected others in a social network, a metric of objective popularity). By extension, the MPFC may also cluster representation of others based on how *subjectively* close we feel to them (i.e., as ‘cliques’ varied by closeness). Another possibility is that interpersonal closeness impinges on our own self-representations, with closer individuals more similarly represented to ourselves. Indeed, social psychology suggests that self-representations include representations of close others (i.e., ‘self-other overlap’) to foster social connection (Aron et al., 1991). Consistent with this possibility, friendship has been associated with sharing similar neural responses to the same social stimuli (Kang et al., 2010; Parkinson et al., 2018), suggesting that interpersonal closeness is tied to self-other similarity. Our first goal was to assess these two possibilities, to determine whether and how MPFC may represent subjective connection between the self and others.

If the MPFC organizes self and other representations in either of these ways, a corollary question is whether it may provide insight into conditions characterized by a chronic lack of subjective social connection, such as loneliness. Loneliness is defined as perceived social isolation, as many lonely individuals maintain multiple social relationships (Berscheid and Reis, 1998; Binder et al., 2012; Jones, 1981; Russell et al., 2012). In fact, the discrepancy between subjective and objective social connection has made loneliness particularly challenging to quantify objectively. Thus, our second goal was to examine if loneliness is associated with altered self-other representation in the MPFC. For example, if the MPFC keeps an organized map of our social circles, loneliness may be associated with alterations in this map. In addition to this possibility, loneliness may be reflected by a lonelier ‘neural self’, with less similarity in representations between the self and others.

To test these possibilities, we assessed neural responses during a self- and other-reflection task, and richly sampled social targets that varied in subjective closeness to participants. We were therefore able to test whether the MPFC 1) keeps an organized map of our interpersonal connections and/or 2) represents the self and others more similarly as a function of subjective closeness, and whether 3) loneliness modulates these patterns.

## Materials and Methods

### Participants

Fifty college students and community members (30 female) between the ages of 18 and 47 (M= 20.2, SD= 4.6) participated in this study. All participants were screened for compliance with MRI safety, reported normal neurological history, and had normal or corrected-to-normal visual acuity. Each participant provided informed consent in accordance with the guidelines set by the Committee for the Protection of Human Subjects at Dartmouth College and received monetary compensation or class credit for participating in the study. FMRI data were excluded for participants (n = 7) whose movement during any run of the scan exceeded 3mm in translation or 2 degrees in rotation. Two additional participants did not complete the Revised UCLA Loneliness Scale (Russell et al., 1980) and were excluded from analyses requiring that measure.

### Procedure

Prior to the scan, participants completed a short survey in which they provided the names of a) five close others with whom they had “the closest, deepest, most involved, and most intimate relationships” and b) five acquaintances (such as classmates, colleagues, or neighbors), ranked in the order in which they felt closest to them. These names were used in the fMRI self- and other-reflection (i.e., trait judgment) task to elicit activation associated with thinking about close others and acquaintances.

During the scan, participants made trait judgments for 16 targets: the self, five nominated close others, five nominated acquaintances, and five well-known celebrities (Ellen Degeneres, Kim Kardashian, Barack Obama, Justin Bieber, and Mark Zuckerberg). Importantly, all targets were familiar to the participants but were expected to vary in subjective closeness. During each 2s trial, one target name was presented above and a trait adjective presented below a central fixation cross. Participants were instructed to consider and respond with how much the trait describes the person (1= “not at all”, 4= “very much”) using a button-box. The task lasted for 10 functional runs (80 trials each, 5 trials per target) for a total of 800 trials (Figure 1).

**Fig. 1.**
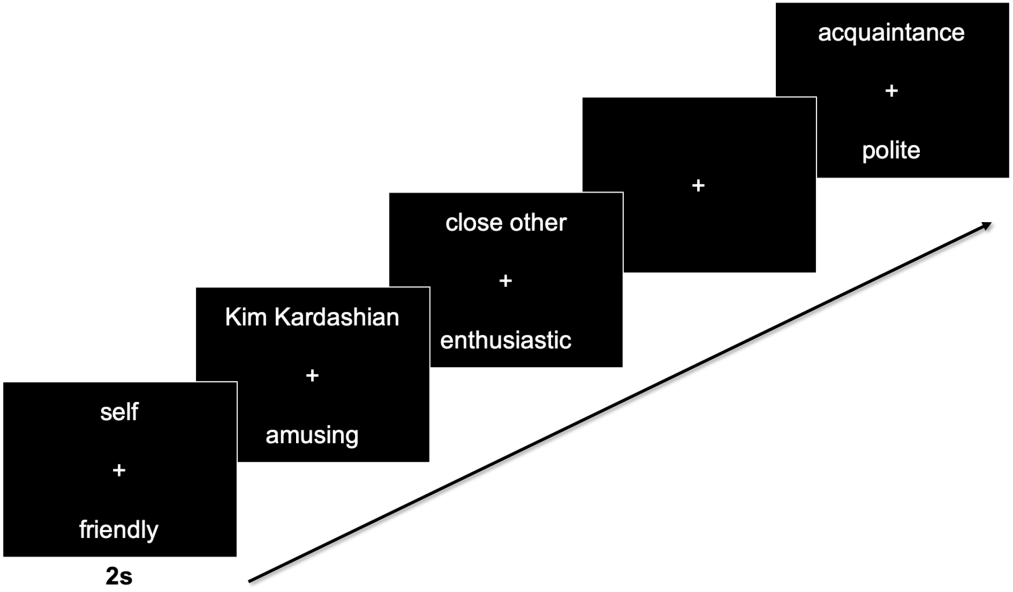
Schematic of self- and other-reflection task design. Participants considered the personality traits of the self, 5 close others, 5 acquaintances, and 5 celebrities across 10 runs. Each trial was presented for 2s and was jittered with fixation.

Following the scan, participants rated their subjective closeness to each of the targets on a 1-100 scale (0= “not at all”, 100= “very much”). Consistent with the idea that people feel closer to their close others, versus acquaintances, versus known celebrities, there was a linear trend in ratings of closeness toward each target, such that close others (M= 82.7, SD= 15.3) were rated closer than acquaintances (M= 46.3, SD= 22.2), who were rated closer than celebrities (M= 7.6, SD= 12.6, β= −53.11, p< 0.001). Afterward, participants also completed the Revised UCLA Loneliness Scale (M= 41.8, SD= 9.4; Russell et al., 1980).

### Apparatus

Imaging data were acquired on a 3T Siemens MAGNETOM Prisma Scanner (Siemens AG, Munich, Germany) with a 32-channel head coil. Stimuli were presented from a 13-inch Apple MacBook Air laptop computer running PsychoPy v1.85 software (Peirce, 2008). An Epson (model ELP-7000) LCD projector displayed the stimuli on a screen at the head end of the scanner bore. Subjects viewed that screen through a mirror mounted on top of the head coil.

### FMRI image acquisition

An anatomical (T1) image was acquired using a high-resolution 3-D magnetization-prepared rapid gradient-echo (MPRAGE) sequence (TR=9.9ms; TE=4.6 ms; flip angle=8°; 1×1×1mm3 voxels). Functional images were collected using a T2*-weighted echo planar imaging (EPI) sequence (TR = 1000 ms, TE = 30ms, flip angle = 59°, bandwidth = 2742, echo spacing = 0.49, 2.5×2.5×2.5 mm resolution) with a simultaneous multi-slice (SMS) of 4 and GeneRalized Auto-calibrating Partial Parallel Acquisition (GRAPPA) of 1. Ten functional runs of 250 axial images (52 slices, 130mm coverage) were acquired for each participant. Sequence optimization was obtained using optseq2 (Dale, 1999) and included 30% jittered trials of fixation for obtaining a baseline estimation of neural activity.

### FMRI preprocessing and response estimation

Univariate functional neuroimaging data were analyzed using SPM12 (Wellcome Department of Cognitive Neurology, London, UK) in conjunction with a suite of preprocessing and analysis tools (https://github.com/wagner-lab/spm12w). Functional data were slice time corrected, realigned within and across runs to correct for head movement and transformed into a standard anatomic space (3-mm isotropic voxels) based on the ICBM 152 brain template space [Montreal Neurological Institute (MNI)]. Normalized data were then smoothed spatially using a 8mm FWHM Gaussian kernel for univariate analyses and a 4mm FWHM Gaussian kernel for multivariate analyses. To further account for motion artifact, participants that demonstrated substantial movement (> 3mm in translation or 2 degrees in rotation) were discarded (n=7, as noted above in the Participants section).

A general linear model (GLM) incorporating task effects (modeled as events of interest convolved with the canonical hemodynamic response function) was used to compute beta images estimating task-related effects for every voxel in the brain. The model included nuisance regressors for six motion parameters (x, y, z directions and roll, pitch, yaw rotations), a linear drift, and run constants. The resulting beta images were used to compute a whole-brain voxel-wise contrast comparing each target to the baseline condition (fixation crosshair trials).

Multivariate analyses were conducted using Python tools for neuroimaging, including the PyMVPA toolkit (http://www.pymvpa.org; Hanke et al., 2009) and SciPy (http://www.scipy.org). Voxel-wise GLM beta images were used to conduct representational similarity analysis (RSA; Kriegeskorte et al., 2008) for the comparison of activation patterns and information overlap across conditions. To interrogate the structure of self-other representation by investigating the (dis)similarity of activation profiles across conditions, a representational (dis)similarity matrix (RDM) was extracted from regions-of-interest (ROIs; see ROI specification section below for more detail). In addition, a sphere searchlight (3mm radius) RSA was conducted to identify regions of the brain that best reflected 1) the (dis)similarity structure identified by the ROI analysis, which distinguished the self, social network members, and celebrities, and 2) self-other closeness ratings. For each searchlight, a one-sample t-test was conducted on the voxel-level Fisher-z transformed correlation values representing similarity of the neural and target RDMs. The resulting group-level statistical maps were voxelwise thresholded at p< 0.001 and cluster-corrected to p<0.001 (minimum extent threshold: k = 195 contiguous voxels for the searchlight analysis of the ROI (dis)similarity structure, k = 157 voxels for the self-other closeness search-light analysis), as recommended by AFNI’s 3dClustSim, using the spatial autocorrelation function (see Eklund et al., 2016; Cox et al., 2017).

### Region-of-Interest (ROI) specification

Given our specific predictions about the role of the MPFC in reflecting connections between the self and others from our social circles, we defined a region-of-interest (ROI) to test the involvement and structure of self-other representation in this region. To define this ROI independently of our own data, we downloaded a reverse-inference map generated by Neurosynth (Yarkoni et al., 2011) reflecting meta-analytic association with the term “self”. Past work suggests that MPFC voxels in Brodmann Area 10 in particular are associated with self-representation (Denny et al., 2012; Wagner et al., 2012; Meyer and Lieberman, 2018; Lieberman et al., 2019). Thus, we further restricted the MPFC ROI to coordinates in BA 10: in the x-dimension from −18 to 18, in the y-dimension from 30 to 80, and in the z-dimension from −12 to 22, for a resulting mask size of 404 voxels (Figure 2A). As shown in the Results section, our whole-brain analyses additionally implicated the posterior cingulate in neural self-other overlap. We therefore defined an *a posteriori* ROI for the posterior cingulate cortex (PCC) using the same approach described for the MPFC, in order to additionally probe how neural self-other overlap in this region relates to loneliness. That is, we used the same meta-analytic map for “self” and further restricted the mask in the x-dimension from −18 to 18, in the y-dimension from −70 to −32, and in the z-dimension from 6 to 44 to isolate the PCC, for a resulting mask size of 223 voxels.

**Fig. 2.**
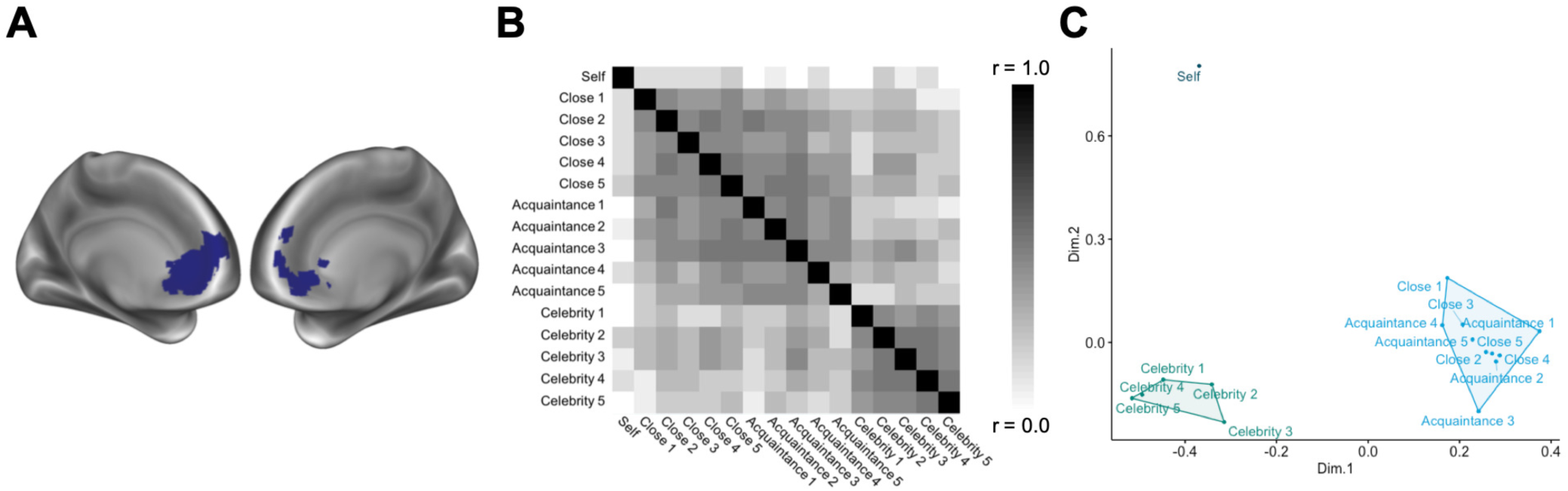
A. ROI mask of the MPFC created from a meta-analytic search for the term “self”, restricted to BA 10 coordinates (Yarkoni et al., 2011). B. Representational similarity of the self and others in the MPFC ROI reflected in a correlation matrix. C. A multi-dimensional scaling plot depicting correlation distances in the MPFC on two explanatory dimensions.

### Analysis of social category representation in MPFC

To examine whether MPFC keeps a structured map of the self and others from our social circles, a neural representational dissimilarity matrix (RDM) was extracted from the MPFC ROI. This approach allowed us to compare the similarity of multivariate activation patterns associated with each target condition. We subsequently compared it with a descriptive, theoretical RDM that appeared to best match the MPFC pattern—distinguishing the self, social network members, and celebrities (see Results). That is, for each subject, the MFPC RDM was correlated with the theoretical RDM, and these individual Fisher z-transformed correlation values were submitted to a group-level t-test. To identify additional brain regions whose response patterns reflected the social category structure identified in the MPFC ROI, a sphere searchlight (3mm radius) was used to conduct a whole-brain RSA with this structure as the model (Figure 3A).

**Fig. 3.**
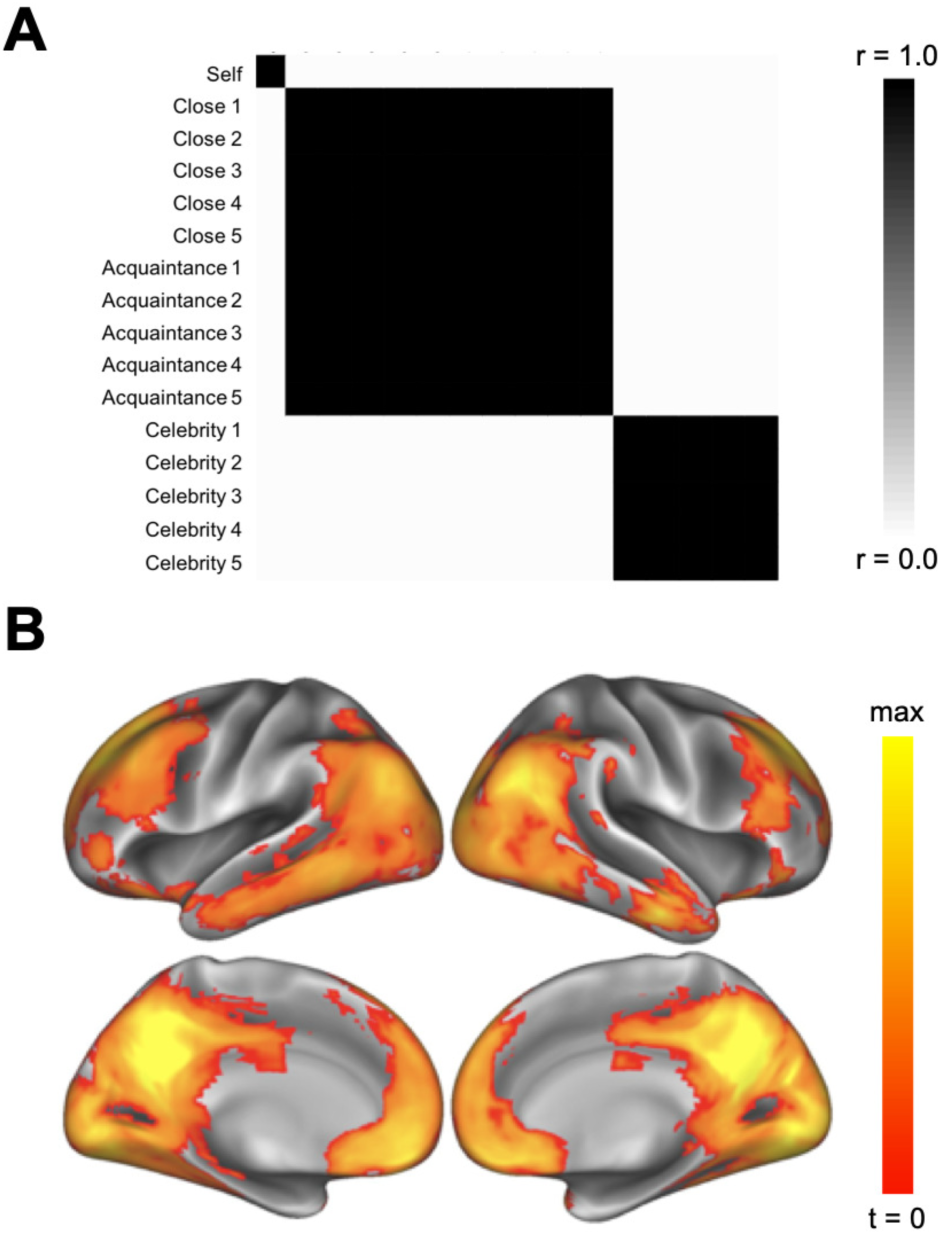
A. Target RDM reflecting a theoretical three-factor structure clustering the self, social network members (close others and acquaintances), and celebrities. B. A whole-brain searchlight RSA revealed brain regions whose similarity structures reflected this target structure (voxel-wise threshold p< 0.001, cluster-corrected p< 0.001).

### Modulation of neural responses by self-other closeness

To determine whether neural responses in the MPFC were sensitive to subjective closeness between the self and others, we performed both univariate and multivariate analyses with the MPFC ROI. We extracted both univariate and multivariate beta estimates from the MPFC ROI and for both: 1) searched for a linear increase in neural response as a function of target condition (self, close others, acquaintances, celebrities) in a linear mixed effects model with a random intercept for subject, and 2) regressed neural responses with self-other closeness ratings for each target in a linear mixed effects model with a random intercept for subject and target, using the *lme4* (Bates et al., 2015) and *lmerTest* (Kuznetsova et al., 2017) packages in R. In addition, we completed follow-up, whole-brain analyses testing for regions across the brain whose activation amplitude and multivariate pattern similarity were modulated by self-other closeness. Reported degrees of freedom for linear mixed effects reflect the total number of subjects contributing to the effect.

#### Univariate Analyses Assessing the Impact of Self-Other Closeness

First, activation betas associated with each target were extracted from the MPFC ROI and were regressed with 1) target condition (self > close others > acquaintances > celebrities) and 2) subjective measures of self-other closeness for each target. Second, a whole-brain parametric modulation analysis was conducted to identify regions of the brain whose activation magnitude increased with self-other closeness. A subject-level regressor reflecting the closeness ratings for each target (from 1-100, with closeness to the self defined as 100) were entered into the first-level GLM to identify brain regions whose activity linearly increased with self-other closeness. Next, a second-level one-sample t-test was conducted and the resulting group-level map was voxelwise thresholded at p< 0.001 and cluster corrected to p< 0.001 (minimum extent threshold: k = 66 contiguous voxels), as recommended by AFNI’s 3dClustSim, using the spatial autocorrelation function (see Eklund et al., 2016; Cox et al., 2017).

#### Multivariate Analyses Assessing the Impact of Self-Other Closeness

Multivariate responses in the MPFC ROI were extracted for a comparison of activation patterns between the self and others, across targets. The similarity (Fisher z-transformed correlation value) between each target and the self were then regressed with 1) target condition (close others > acquaintances > celebrities) and 2) self-other closeness ratings for each target. To determine whether additional brain regions demonstrate increases in neural similarity between the self and others with greater closeness, a sphere searchlight (3mm radius) was used to conduct a whole-brain RSA. Specifically, this model assessed similarity in multivariate responses between the self condition and each other target, weighted by the subject’s closeness rating for that target (i.e., comparing neural patterns to the self with each target, weighted by each subject’s vector of 15 closeness ratings). The 1-100 closeness ratings were transformed to a 0-1 scale and converted to distances (1 minus the transformed value) to permit correlation with neural representational dissimilarity matrices. Similarity in activation patterns was estimated using Pearson correlation and similarity across brain activation patterns and behavioral responses were estimated using Spearman rank correlation. These correlation values were Fisher z-transformed to allow for statistical comparisons between conditions.

### Relating self-other neural responses in MPFC and PCC to loneliness

Next, we sought to determine whether loneliness alters the representation of subjective social connections in MPFC, as well as PCC. We examined two possibilities: 1) whether loneliness alters the mapping of social circles in MPFC and PCC and 2) whether loneliness is associated with greater dissimilarity in representations between the self and others in MPFC and PCC. To test the first possibility that loneliness may be associated with altered mapping of social circles, we extracted pairwise neural similarity (Fisher-z transformed correlation coefficients) in each of these ROIs to examine within-condition similarity for each social circle (i.e., close others, acquaintances, and celebrities) and between-condition similarity as a function of the distance between the social circles, according to their closeness to the self (i.e., at each step from the diagonal of the RDM). This revealed neural similarity for targets within a condition (e.g. how similarly close others are represented to one another, how similarly acquaintances are represented to one another, and how similarly celebrities are represented to one another; distance of 0), from adjacent conditions (e.g., close others and acquaintances, and separately, acquaintances and celebrities; distance of 1), and from distant conditions (close others and celebrities; distance of 2). In these models, pairwise neural similarity among targets was predicted by both loneliness and either 1) the distance between target conditions (or social circles) in the RDM or 2) the particular target condition (for targets in the same social circle). A median split on loneliness (median = 43) was conducted for visualization and for examining simple effects. All other models used continuous loneliness values. Full models predicting similarity from loneliness and target condition or between-condition distance included a random intercept for subject. To avoid overfitting, models testing simple effects of loneliness on neural similarity for a particular target condition or between-condition distance included a random intercept for the particular target pair (e.g., close 1 – close 2).

Finally, to test the second possibility that loneliness is associated with greater dissimilarity in representations between the self and others in MPFC and PCC, participants’ loneliness scores were entered as a predictor of univariate neural activation and multivariate self-other neural similarity (Fisher-z transformed correlation coefficients of each condition with the self) in each of these ROIs, along with either target condition or self-other closeness ratings. Specifically, separate regression models predicted activation magnitude in either of these regions as a function of: 1) target condition, linearly ordered by closeness to the self (self > close others > acquaintances > celebrities) or 2) subjective self-other closeness ratings to each target. Similarly, separate regression models predicted self-other multivariate similarity in these regions as a function of: 1) target condition (close others > acquaintances > celebrities) or 2) subjective self-other closeness ratings. The models with self-other closeness as a predictor included a random intercept for target; to avoid overfitting, those with target condition as a predictor did not.

## Results

### Does MPFC maintain an organized map of ourselves and our social circles?

#### Social category representation in MPFC

To interrogate the structure of self-other representation in the MPFC, we extracted the neural representational dissimilarity matrix (RDM) from the MPFC ROI (Figure 2A). The overall representational structure in the MPFC appeared to differentiate the self from all other conditions while the representation of others was clustered such that all social network members (close others and acquaintances) were similarly represented to one another and celebrities were, separately, similarly represented to one another (Figure 2B). To better visualize the emergent structure of the matrix, a multi-dimensional scaling solution was derived to depict the similarity of conditions along two dimensions (Figure 2C). This plot more clearly illustrates that three clusters emerge from cross-condition similarity in MPFC activation: self, social network members (close others and acquaintances) and celebrities. In fact, the neural RDM from this region correlated with this social category structure, r = 0.34, t(42) = 10.78, p< 0.001. These results provide evidence that the MPFC preserves information about closeness to the self in a categorical, clique structure—but it may do so by collapsing social network members (i.e. close others and acquaintances).

#### Social category representation across the whole-brain

To search for other brain regions with representational profiles similar to the one reflected in the MPFC, a whole-brain searchlight analysis was performed using the three-cluster, social category structure as the target RDM (Figure 3A). This analysis yielded regions across the social brain, voxel-wise threshold p< 0.001, cluster-corrected p< 0.001 (Table 1). In addition to MPFC, this analysis revealed a cluster in PCC/precuneus that extended into temporoparietal junction, middle temporal gyrus, and temporal poles (Figure 3B). Here we demonstrate that the MPFC crudely organizes representations of individuals from our social circles according to their closeness, whereas the self remains distinct. Moreover, this structure was observed throughout the social brain.

**Table 1.**
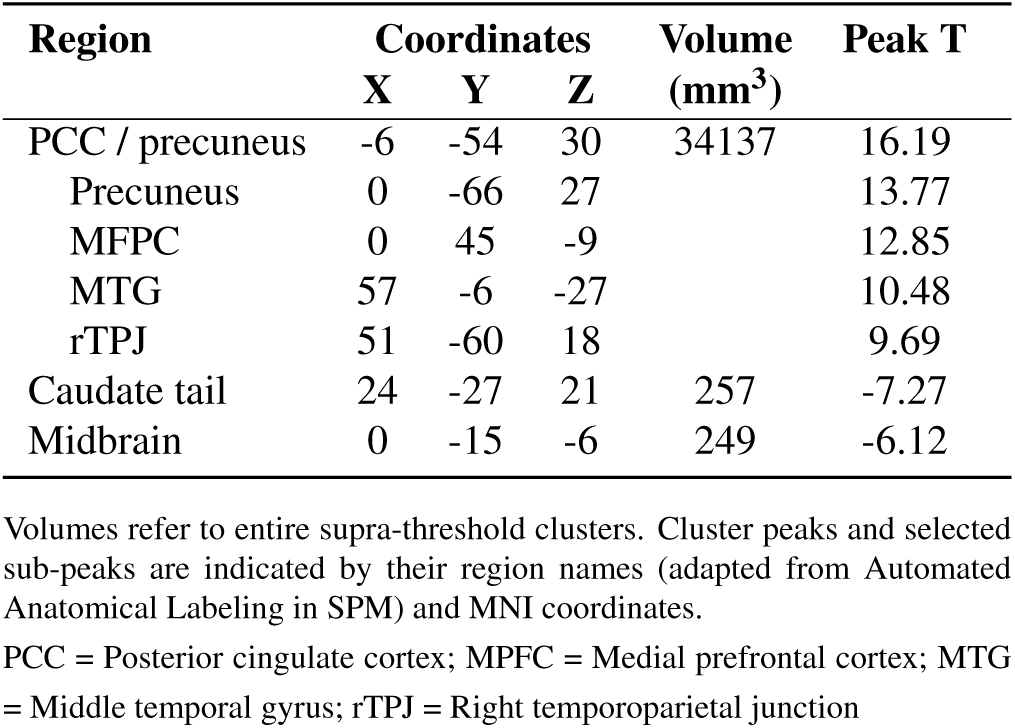
Peak regions from searchlight analysis for three-factor structure clustering the self, social network members (close others and acquaintances), and celebrities (voxel-wise threshold p< 0.001, cluster-corrected p< 0.001).

| Region | Coordinates |  |  | Volume (mm<sup>3</sup>) | Peak T |
| --- | --- | --- | --- | --- | --- |
|  | X | Y | Z |  |  |
| PCC / precuneus | -6 | -54 | 30 | 34137 | 16.19 |
| Precuneus | 0 | -66 | 27 |  | 13.77 |
| MPFC | 0 | 45 | -9 |  | 12.85 |
| MTG | 57 | -6 | -27 |  | 10.48 |
| rTPJ | 51 | -60 | 18 |  | 9.69 |
| Caudate tail | 24 | -27 | 21 | 257 | -7.27 |
| Midbrain | 0 | -15 | -6 | 249 | -6.12 |

Volumes refer to entire supra-threshold clusters. Cluster peaks and selected sub-peaks are indicated by their region names (adapted from Automated Anatomical Labeling in SPM) and MNI coordinates.
PCC = Posterior cingulate cortex; MPFC = Medial prefrontal cortex; MTG = Middle temporal gyrus; rTPJ = Right temporoparietal junction

### Does subjective self-other closeness modulate similarity in MPFC responses to self and others?

#### Neural responses to self-other closeness in MPFC

To isolate the influence of social closeness on neural responses to the self and others, we began by assessing whether univariate response magnitudes in the MPFC were sensitive to the participant’s closeness to each target. Consistent with this prediction, activation magnitudes in the *a priori*-defined MPFC ROI linearly increased with the social closeness of the target to the participant (linear effect by target condition: β= 0.33, t(43)= 8.59, p< 0.001; linear increase with social 1.43, p= 0.15), and only marginally increased as a func tion of target-specific self-other closeness ratings (β= 0.02, t(43)= 1.96, p= 0.066). However, we found evidence for greater neural self-other overlap with close others compared to acquaintances and celebrities. That is, the direct comparison of neural self-other overlap with close others relative to both acquaintances and celebrities was significant (β= 0.06, t(43)= 2.70, p= 0.007; Figure 5). These findings provide novel and rigorous support for the social psychological idea that close others are incorporated into our own self-representations (Aron et al., 1991).

#### Searchlight analysis for self-other closeness

For a more extensive exploration of how social closeness moduates self-other neural similarity, we conducted a whole-brain searchlight analysis on self-other closeness ratings. That is, closeness ratings were used as the comparison metric (i.e., target similarity metric) for self-other neural similarity. This analysis revealed regions where targets who are personally close to the self also elicit similar activation patterns as the self. This searchlight analysis revealed regions across the social brain, including the PCC/precuneus and MPFC, whose patterns of self-other similarity best matched the sub-closeness ratings: β= 0.13, t(43)= 5.67, p< 0.001, Figure 4A). This result complements and extends previous literature demonstrating intermediate activation of the MPFC for a single close other relative to the self and a single distant other (Feng et al., 2018; Krienen et al., 2010).

**Fig. 4.**
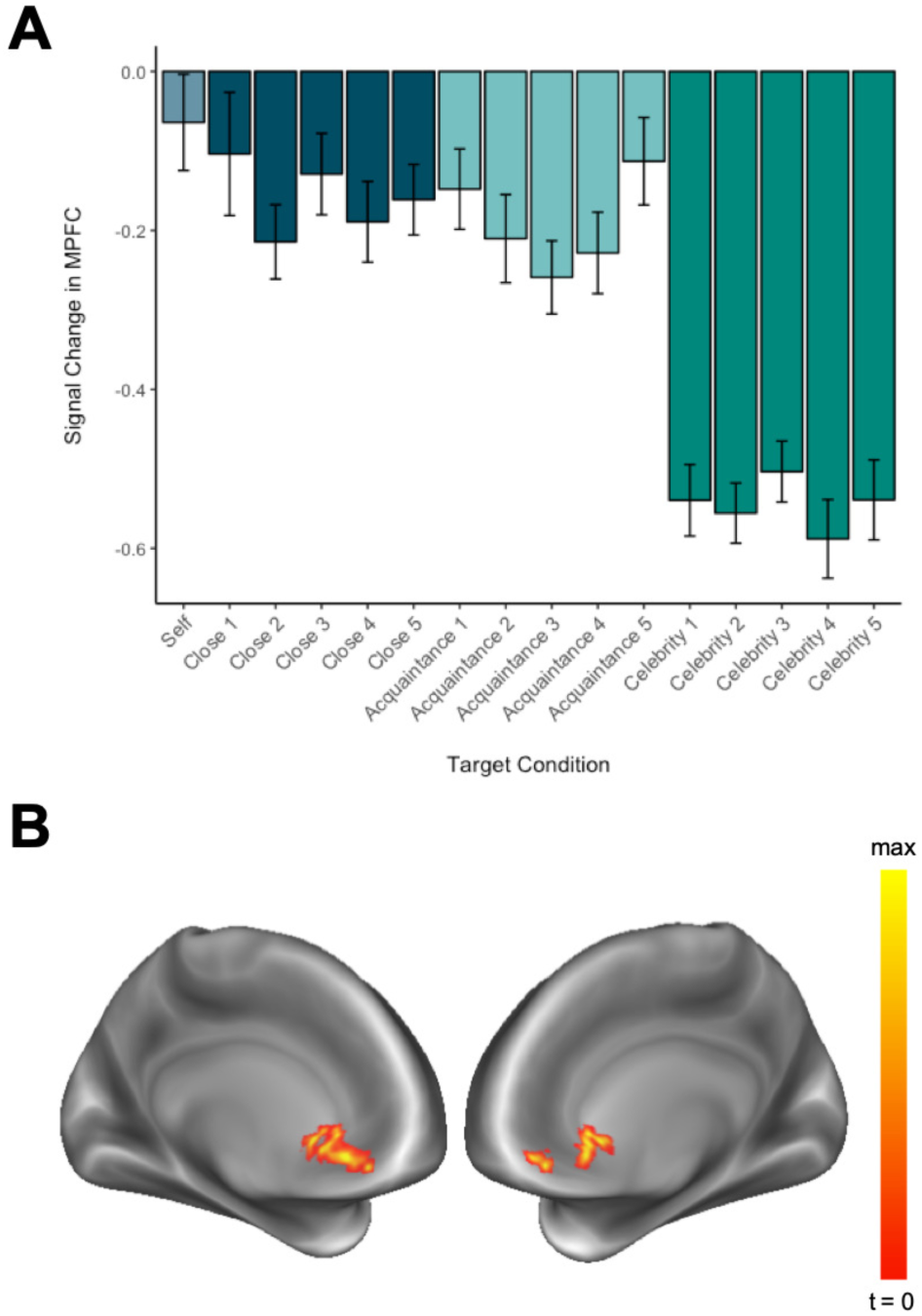
A. Activation magnitude in the MPFC ROI linearly increased with self-other closeness (linear effect by target condition: β= 0.33, t(43)= 8.59, p< 0.001; linear increase with subjective closeness ratings: β= 0.13, t(43)= 5.67, p< 0.001). B. Whole-brain parametric modulation analysis revealed a single cluster in the MPFC (−3, 33, −9) whose activation magnitude linearly increased with the self-other closeness of the participant and target (voxelwise p< 0.001, cluster-corrected to p< 0.001).

#### Parametric modulation by self-other closeness across whole brain

Next, we conducted a whole-brain parametric modulation analysis to more extensively search for brain regions that were sensitive to the closeness of the target considered during the task. This analysis revealed a single cluster in the MPFC (MNI: −3, 33, −9; voxelwise p< 0.001, cluster-corrected to p< 0.001) whose BOLD activation linearly increased with the social closeness of the target to the participant (Figure 4B). Both univariate analyses suggested that activation in the MPFC, defined *a priori* for its association with self-related processing and identified whole-brain, is modulated by social closeness.

#### Neural representation of self-other closeness in MPFC

To determine whether multivariate activation patterns in the MPFC were sensitive to subjective self-other closeness, we extracted parameter estimates that reflected the similarity of each target to the self in this region. Self-other overlap did not linearly increase with target condition (β= 0.03, t(43)= 1.43, p= 0.15), and only marginally increased as a function of target-specific self-other closeness ratings (β= 0.02, t(43)= 1.96, p= 0.066). However, we found evidence for greater neural self-other overlap with close others compared to acquaintances and celebrities. That is, the direct comparison of neural self-other overlap with close others relative to both acquaintances and celebrities was significant (β= 0.06, t(43)= 2.70, p= 0.007; Figure 5). These findings provide novel and rigorous support for the social psychological idea that close others are incorporated into our own self-representations (Aron et al., 1991).

**Fig. 5.**
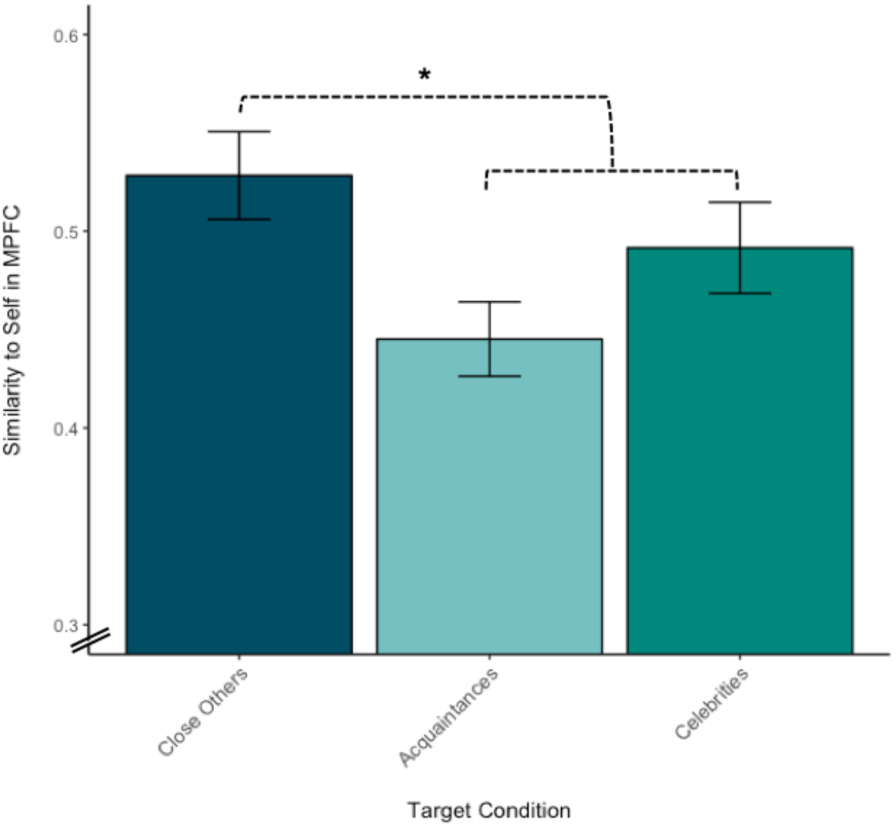
Self-other neural similarity was greater for close others than for other targets (acquaintances and celebrities) in the MPFC, β= 0.06, t(43)= 2.70, p= 0.007.

### Searchlight analysis for self-other closeness

For a more extensive exploration of how social closeness moduates self-other neural similarity, we conducted a whole-brain searchlight analysis on self-other closeness ratings. That is, closeness ratings were used as the comparison metric (i.e., target similarity metric) for self-other neural similarity. This analysis revealed regions where targets who are personally close to the self also elicit similar activation patterns as the self. This searchlight analysis revealed regions across the social brain, including the PCC/precuneus and MPFC, whose patterns of self-other similarity best matched the subjective closeness ratings reported by the participants (voxelwise threshold p< 0.001, cluster-corrected p< 0.001; Table 2; Figure 6).

**Table 2.** Peak regions from searchlight analysis comparing self-other neural similarity to social closeness ratings, voxel-wise threshold p< 0.001, cluster-corrected p< 0.001.

| Region | Coordinates |  |  | Volume (mm<sup>3</sup>) | Peak T |
| --- | --- | --- | --- | --- | --- |
|  | X | Y | Z |  |  |
| MPFC | 9 | 51 | 33 | 235 | 6.43 |
| Dorsal MPFC | 12 | 39 | 45 |  | 5.27 |
| Right SFG | 21 | 33 | 42 |  | 3.93 |
| PCC / Precuneus | -6 | -24 | 39 | 659 | 6.51 |
| Mid-cingulate | -3 | -15 | 39 |  | 6.27 |
| Mid-cingulate | 3 | -24 | 42 |  | 6.04 |
| Visual cortex | -6 | -90 | -3 | 2957 | 12.34 |
| Lingual gyrus | 6 | -84 | -3 |  | 11.67 |
| Lingual gyrus | 15 | -84 | -6 |  | 11.61 |
| Left SFG | -24 | 36 | 45 | 260 | 6.40 |
| Left SFC | -24 | 27 | 42 |  | 5.31 |
| Left MFG | -33 | 24 | 48 |  | 4.91 |

Volumes refer to entire supra-threshold clusters. Cluster peaks and selected sub-peaks are indicated by their region names (adapted from Automated Anatomical Labeling in SPM) and MNI coordinates.
MPFC = Medial prefrontal cortex; PCC = Posterior cingulate cortex; SFG = Superior frontal gyrus; SFC = Superior frontal sulcus; MFG = Middle frontal gyrus

**Fig. 6.**
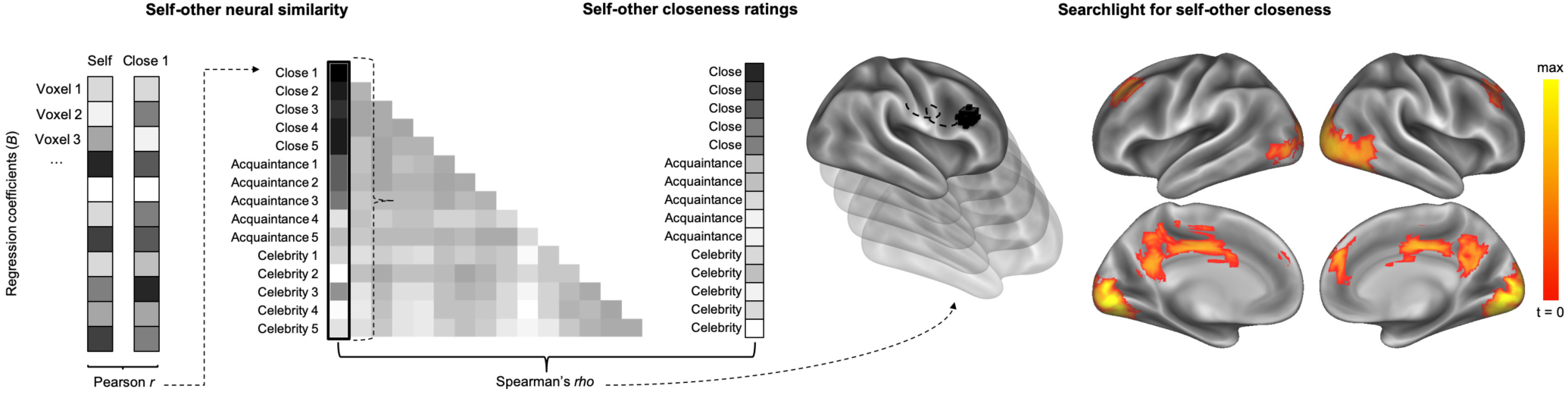
A whole-brain searchlight representational similarity analysis (RSA) for self-other closeness. Brain regions where self-other neural similarity was positively correlated with ratings of self-other closeness included the PCC/precuneus and MPFC (voxel-wise threshold p< 0.001, cluster-corrected p< 0.001).

Overall, it appears that the magnitude of neural activity and multivariate representation in MPFC in response to other targets become increasingly similar to the self with subjective social closeness.

### Is loneliness associated with altered responses to the self and others in MPFC and PCC?

To test the possibility that neural responses to the self and others could reflect meaningful individual differences in social connection, we related both univariate and multivariate parameter estimates in the MPFC and PCC ROIs to trait loneliness. These regions were selected for their association with both social category and self-other closeness representation in the previous whole-brain analyses. These ROIs were defined independently of our own data (see Methods) for the following analyses.

#### Loneliness modulates univariate activation to the self and others in MPFC

Examining neural activation magnitude in MPFC across all target conditions revealed a main effect (ME) of loneliness whereby greater loneliness was associated with less MPFC activation (β= −0.065, t(41)= −3.43, p< 0.001); this effect did not differ across target conditions (interaction between linear trend for condition and loneliness: β= 0.055, t(41)= 1.24, p= 0.22). In a similar model of MPFC activation by self-other closeness and loneliness, there was a main effect of loneliness whereby greater loneliness was associated with less MPFC activation (β= −0.032, t(41)= −2.12, p= 0.035); this effect did not differ as a function of self-other closeness (interaction of self-other closeness and loneliness: β= 0.016, t(41)= 1.02, p= 0.31). That is, lonelier individuals may exhibit less MPFC activity while reflecting on people, regardless of the person considered. In contrast, loneliness was not associated with neural activation in the PCC (ME loneliness in model with condition: β= −0.011, t(41)= −0.53, p= 0.59; ME loneliness in model with self-other closeness: β= −0.003, t(41)= −0.19, p= 0.85) and did not differ across conditions (β= 0.01, t(41)= 0.19, p= 0.85) or as a function of self-other closeness (β= 0.014, t(41)= 0.78, p= 0.44).

#### Loneliness modulates representation of social circles in MPFC and PCC

To test whether loneliness modulates the mapping of social circles in MPFC and PCC, we first compared neural (dis)similarity of each pair of targets, excluding the self condition, as a function of their distance from the self (i.e., at each step from the diagonal of the similarity matrix). This analysis assesses how similarly targets within a condition are represented (distance of 0), how similarly targets from adjacent conditions are represented to one another (close others and acquaintances, and separately, acquaintances and celebrities (distance of 1), and how similarly the two most distant target conditions, close others and celebrities, are represented to one another (distance of 2).

Overall, we observed a decay in the neural similarity of social targets with increasing social distance between the social circles in MPFC and PCC. Within condition similarity was high (distance of 0), but neural similarity decreased as the distance between conditions increased—both linearly and quadratically in the MPFC (linear effect: β= −0.21, t(41)= - 4.49, p< 0.001, quadratic effect: β= 0.11, t(41)= 2.85, p= 0.004; Figure 7A) and quadratically in the PCC (linear effect: β= −0.064, t(41)= −1.38, p= 0.17, quadratic effect: β= 0.11, t(41)= 2.71, p= 0.007; Figure 7B).

**Fig. 7.**
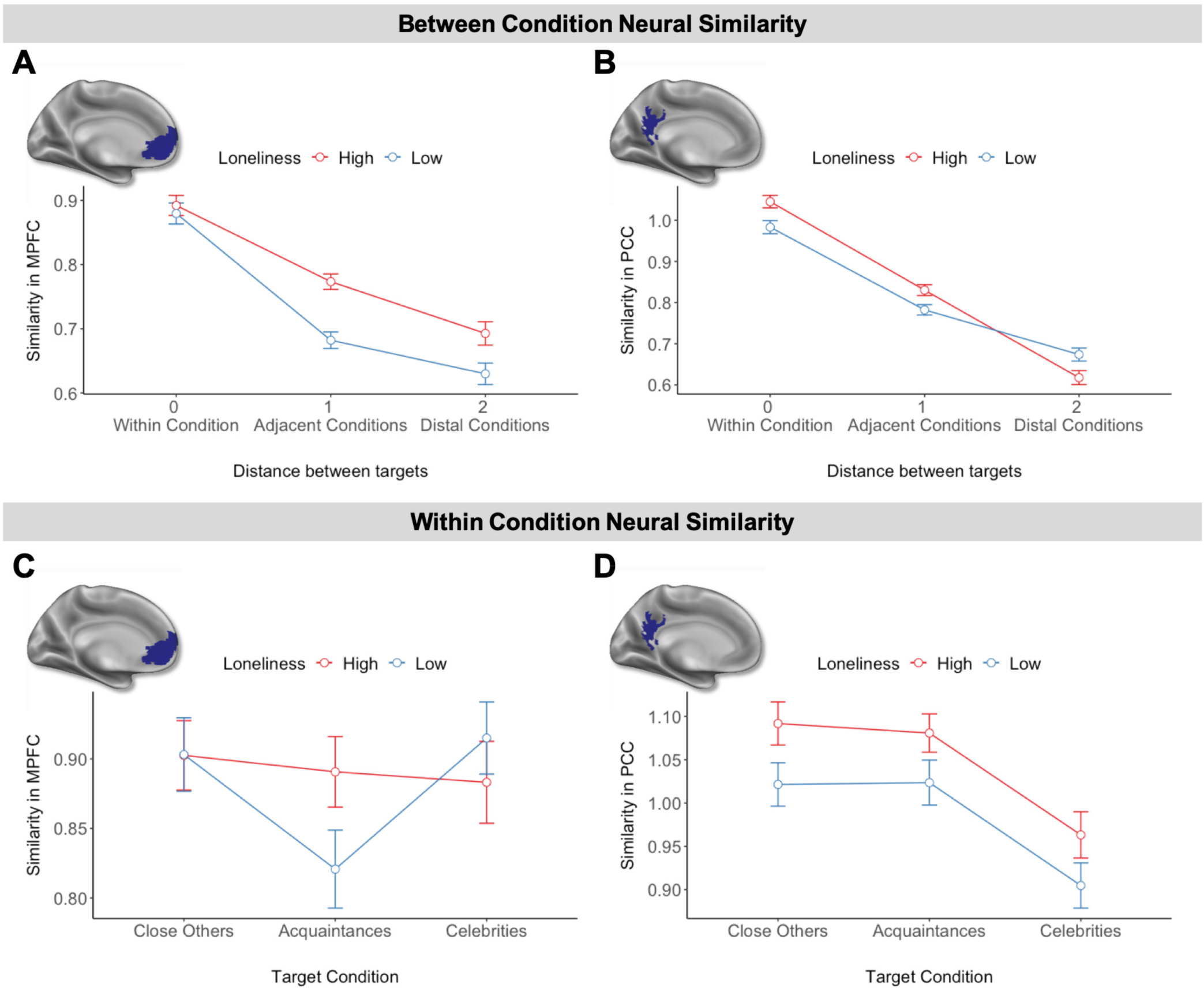
A. Between condition neural similarity in MPFC decreased with the distance between social categories relative to the self. This trend was both linear and quadratic (i.e., leveling off at greater distances between target conditions) for less lonely people, but it decreased linearly for lonelier people. B. Similarly, between condition neural similarity in PCC decreased with social distance, and the pattern was both linear and quadratic for less lonely people but linear for lonelier people. C. Within condition neural similarity in MPFC for high and low lonely individuals. D. Within condition neural similarity in PCC for high and low lonely individuals. Loneliness data were median split for visualization but used continuously during analysis.

Critically, loneliness moderated the decay in similarity across these social circles. Loneliness increased the linearity of the decay in PCC (β= −0.005, t(41)= −4.32, p< 0.001) and decreased the degree of quadratic decay in PCC (β= - 0.002, t(41)= −2.28, p= 0.023) and marginally in MPFC (β= −0.002, t(41)= −1.95, p= 0.051). For those low in loneliness, the decay in social target similarity leveled off at greater social distances from the self, reflecting both a linear (MPFC: β= −0.18, t(20)= −12.65, p< 0.001; PCC: β= −0.22, t(20)= - 16.11, p< 0.001) and quadratic trend (MPFC: β= 0.06, t(20)= 5.08, p< 0.001; PCC: β= 0.04, t(20)= 3.33, p< 0.001). In contrast, lonelier participants showed a stronger linear trend (MPFC: β= −0.14, t(21)= −10.35, p< 0.001; PCC: β= −0.31, t(21)= −21.66, p< 0.001), and no quadratic trend (MPFC: β= 0.02, t(21)= 1.37, p= 0.17; PCC: β= 0.0009, t(21)= 0.08, p= 0.96). More specifically, as loneliness increased, adjacent conditions (i.e., close other and acquaintances; acquaintances and celebrities; distance of 1) were represented more similarly to one another in both the MPFC (β= 0.004, t(41)= 4.87, p< 0.001) and PCC (β= 0.007, t(41)= 7.91, p< 0.001; Figure 7A and 7B). In MPFC, this blurring of social circles with loneliness was even observed in the similarity between close others and celebrities (i.e., distance of 2; β= 0.003, t(41)= 2.40, p= 0.017, Figure 7A).

In addition to changes *between* social circles, loneliness may alter *within* social circle mapping as well. To explore this possibility, we examined pairwise neural similarity for social targets belonging to the same social circle. Most strikingly, as loneliness increased, acquaintances were represented more similarly to one another in the MPFC (β= 0.004, t(41)= 1.98, p= 0.049, Figure 7C) and PCC (β= 0.008, t(41)= 3.96, p< 0.001, Figure 7D). Close others (β= 0.008, t(41)= 3.73, p<= 0.001) and celebrities (β= 0.008, t(41)= 3.85, p<= 0.001) were also represented more similarly to one another in PCC (Figure 7D).

Collectively, these results suggest that loneliness is associated with an altered neural map of social circles. Although there were some differences in this alteration between MPFC and PCC, in both of these regions loneliness was associated with blurred boundaries *between* the social circles surrounding acquaintances (i.e., increased similarity between close others and acquaintances, as well as acquaintances and celebrities), and blurred boundaries among the collection of acquaintances themselves (i.e., increased similarity between acquaintances).

#### Loneliness modulates self-other similarity in MPFC and PCC

Finally, to test the possibility that loneliness is associated with a disconnected neural self-representation, we regressed self-other similarity in multivariate activation patterns with the loneliness of the participant separately for the MPFC and PCC ROIs. In the MPFC, loneliness negatively related to self-other similarity across all target conditions (ME loneliness in model with condition: β= −0.05, t(41)= −3.91, p< 0.001; ME loneliness in model with self-other closeness: β= −0.05, t(41)= −3.67, p< 0.001); and this did not differ as a function of target condition (β= 0.008, t(41)= 0.33, p= 0.74; Figure 8A) or self-other closeness (β= −0.003, t(41)= −0.19, p= 0.85). Conversely, in the PCC, loneliness positively related to self-other similarity across all target conditions (ME loneliness in model with condition: β= 0.073, t(41)= 5.17, p< 0.001; ME loneliness in model with self-other closeness: β= 0.08, t(41)= 5.46, p< 0.001); and this did not differ as a function of target condition (β= −0.02, t(41)= −0.82, p= 0.41; Figure 8B) or self-other closeness (β= −0.002, t(41)= −0.13, p= 0.90). These results suggest that lonelier individuals may indeed represent others as more distant or dissimilar from the self in the MPFC. In contrast, lonelier individuals may represent their self as more connected or similar to others in the PCC.

**Fig. 8.**
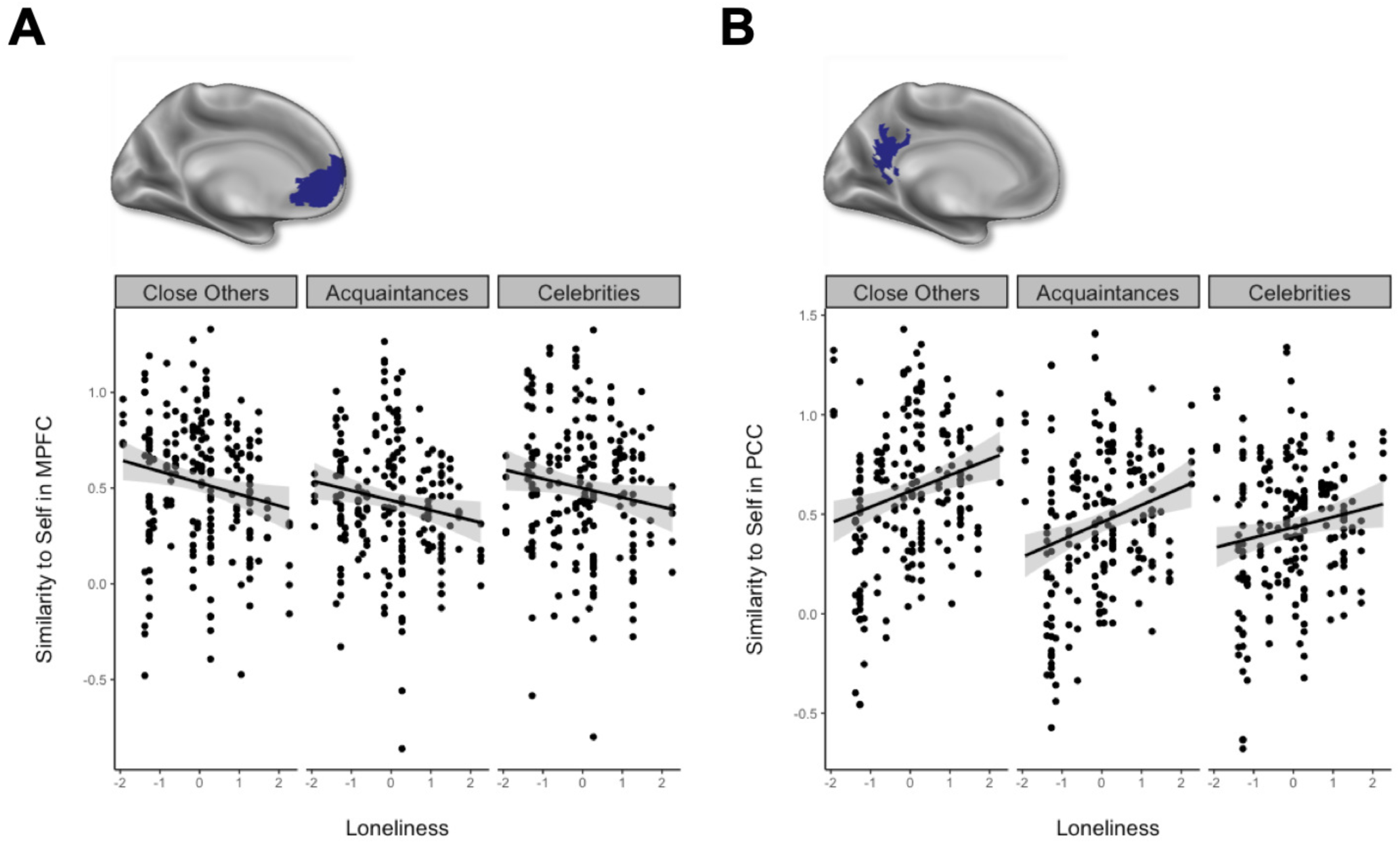
A. Loneliness was associated with lower self-other similarity in the MPFC across all target conditions (β= −0.05, t(41)= −3.91, p< 0.001). B. Conversely, loneliness was associated with greater self-other similarity in the PCC across all target conditions (β= 0.073, t(41)= 5.17, p< 0.001).

## Discussion

How does the brain represent our subjective social connection to others? Here, we found that multivariate activation patterns in the MPFC and regions across the social brain reflected a crude categorization of social targets. Specifically, the social brain 1) represents the self distinctly from others and 2) clusters representation of others based on whether or not they are a part of our social network. Although the self was represented distinctly from others, self-other closeness modulated this distance. Greater subjective self-other closeness was associated not only with greater response amplitudes in MPFC, but also greater overlapping multivariate patterns of neural activity in both MPFC and throughout the social brain. Critically, loneliness moderated the organization of the self and others, with blurred representations of weaker ties (i.e. acquaintances), and altered neural self-other overlap. Collectively, these results provide novel insight into how the social brain maps our subjective connections to others, and how loneliness alters this mapping.

Most people maintain a core set of 3-5 close others but regularly interact with larger groups of friends and acquaintances, with diminishing frequency and emotional attachment for more peripheral relationships (Zhou et al., 2005; Roberts and Dunbar, 2011; Dunbar, 2018). Indeed, it is thought that the time, effort, financial, and even information processing costs of maintaining relationships constrain the size of each of these social network layers (Stiller and Dunbar, 2007; Roberts and Dunbar, 2011). Thus, both the broad clustering of individuals based on whether or not they are in our social network (close others and acquaintances vs. celebrities) and the more fine-grained increase in self-other overlap with greater interpersonal closeness may help us efficiently respond to individuals in our social environment. To further test this possibility, future work may examine whether individuals with the largest social networks also have the most efficiently organized representations between the self and others. Just as clearer physical maps help us navigate physical space, clearer interpersonal maps may help us navigate social interactions.

### Self-other closeness increases neural representational overlap

To our knowledge, our results provide the most rigorous assessment to date of increasing neural self-other overlap with interpersonal closeness, finding evidence for self-other overlap in both MPFC and PCC. Self-other overlap is an important construct in social psychology: it is a defining feature of interpersonal relationships (Aron et al., 1991; Branand et al., 2019) and corresponds with pro-social outcomes, such as enhanced empathy (Galinsky et al., 2005). Yet, it has been difficult to precisely test for overlapping representations between the self and others, in part due to methodological limitations. Self-other overlap is typically assessed by 1) asking participants to indicate the degree of overlap they feel they share with others (Aron et al., 1991), 2) source errors between the self and others in memory (Benoit et al., 2010; Bergström et al., 2015), and/or 3) comparing univariate activity in MPFC to the self and a single close other and non-close other (Krienen et al., 2010). Given that none of these methods assess representational content, evidence for self-other overlap from these measures cannot rule out alternative interpretations. We capitalized on multivariate neural pattern similarity to more precisely demonstrate overlapping representations between the self and others as a function of closeness. Building on prior literature (Schmitz and Johnson, 2007), we suggest that the blurring of self-other representations observed here could be due to the interpersonal nature of the self, the richness with which personally known others (including the self) are represented, and/or the heightened subjective value associated with relationship partners.

For example, some have suggested that the self can only be defined in the context of its relationship to others (Andersen and Chen, 2002). As such, a latent, relational version of the self may exist for each unique relationship, which comprises all aspects of the self that are most relevant for that relationship (Andersen and Chen, 2002). Thus, neural self-other overlap may reflect the relational self associated with the target considered. Alternatively, the increase in neural self-other similarity with subjective closeness may reflect more shared experience with these individuals, as a shared history might provide more information to enrich mental models of the person (Schmitz and Johnson, 2007; Murray et al., 2012). Consistent with this possibility, the more experience we have with a person, the richer our multivariate MPFC representation of their mental states (Thornton et al., 2019).

Additionally, social network members may be endowed with enhanced subjective value, which is frequently associated with the ventral MPFC (Wagner et al., 2019). Indeed, the self and subjective value may be intrinsically related (Tamir and Mitchell, 2012; Chavez et al., 2017) and associated with closeness to the self (Schmitz and Johnson, 2007; Murray et al., 2012). Interestingly, the parametric modulation analysis testing for neural activity increasing with closeness to the self yielded a ventral portion of the MPFC similarly located to clusters associated with subjective value (Bartra et al., 2013; Clithero and Rangel, 2014). In contrast, the cluster that emerged in the multivariate self-other overlap analysis was more dorsal, and is a region frequently associated with person perception and social inference (Lieberman et al., 2019). Mean levels of neural activity in vMPFC may track the value of others, whereas shared multivariate responses in more dorsal portions of MPFC may reflect shared representational content between the self and others.

It is noteworthy that the PCC emerged in the self-other overlap analysis, in addition to MPFC. While both regions are associated with self-other distinction (Van Overwalle, 2009; Feng et al., 2018), the MPFC is associated with responding to self > other and the PCC is often associated with responding to other > self (Murray et al., 2015) and might facilitate other-focused social cognition (Johnson et al., 2006). The MPFC and PCC comprise the default network’s core subsystem (Andrews-Hanna et al., 2010; Yeo et al., 2011), meaning that their neural activity often oscillates in conjunction with one another. Our schemas of the self and others may be spread through this network and while overlapping representations in MPFC may reflect the inclusion of the other in the self, overlapping PCC representations may reflect the inclusion of the self in the other.

### Loneliness alters neural self-other representation

Critically, trait-level social connection (i.e. loneliness) modulated self-other representation. Lonelier participants had blurred boundaries in MPFC and PCC between the social circles surrounding acquaintances, as well as blurred boundaries among the collection of acquaintances themselves. Chronic social disconnection may therefore suppress the distinctiveness with which peripheral social network members are represented. Interestingly, social network research emphasizes the ‘strength of weak ties,’ specifically highlighting the important role acquaintances play in well-being, social support, and access to information (Granovetter, 1973; Wellman and Wortley, 1990; Sandstrom and Dunn, 2014). However, neuroscience research on loneliness focuses on lonely individuals’ responses to either strangers (Cacioppo et al., 2009; Yamada and Decety, 2009) or close others (Inagaki et al., 2016), but not acquaintances. Our findings suggest that understanding how the brain represents weak ties may yield important insight into how loneliness links to negative outcomes.

Loneliness was also associated with self-other similarity in the MPFC and PCC. Specifically, lonelier people represent the self more dissimilarly from others in the MPFC and more similarly to others in the PCC. As mentioned above, the MPFC is associated with responding to self > other, while the PCC/precuneus is often associated with responding to other > self (Murray et al., 2015). In light of these associations, lonely individuals may increasingly view others as more distant from the self and may simultaneously fail to divorce themselves from their representations of others. Intriguingly, the observation that loneliness was associated with more representational distance from others in the MPFC aligns with the phenomenological experience of loneliness. Self-report measures of loneliness point to a disconnected self, with lonelier individuals endorsing statements such as ‘I feel isolated from others’ and ‘people are around me but not with me’ (Russell et al., 1980). Our results suggest that the subjective experience of loneliness can be traced to a lonelier ‘neural self’, with lonelier individuals distancing themselves from their social connections even at the level of neural representation.

## Conclusion

The quality and intimacy of social relationships are critical predictors of happiness and well-being (Klinger, 1977; Diener and Seligman, 2002; Holt-Lunstad et al., 2010). Our results suggest that the social brain may help us navigate our social connections by mapping people based on whether or not they are in our social network, with our closest social ties represented most closely to ourselves. Moreover, loneliness is associated with distortions in this mapping, particularly blurred representations of weak ties and skewed neural similarity between the self and others. The paths we take in social life may depend, in part, on the interpersonal maps we carry in our social brains.

## ACKNOWLEDGEMENTS

Research reported in this publication was supported by the National Institute Of Mental Health of the National Institutes of Health under Award Number F31MH111192. The content is solely the responsibility of the authors and does not necessarily represent the official views of the National Institutes of Health.

